# DNA motifs are not general predictors of recombination in two *Drosophila* sister species

**DOI:** 10.1101/453639

**Authors:** James M. Howie, Rupert Mazzucco, Thomas Taus, Viola Nolte, Christian Schlötterer

## Abstract

Meiotic recombination is crucial for chromosomal segregation, and facilitates the spread of beneficial and removal of deleterious mutations. Recombination rates frequently vary along chromosomes and *Drosophila melanogaster* exhibits a remarkable pattern. Recombination rates gradually decrease towards centromeres and telomeres, with dramatic impact on levels of variation in natural populations. Two close sister species, *D. simulans* and *D. mauritiana* do not only have higher recombination rates, but also exhibit a much more homogeneous recombination rate that only drops sharply close to centromeres and telomeres. Because certain sequence motifs are associated with recombination rate variation in *D. melanogaster*, we tested whether the difference in recombination landscape between *D. melanogaster* and *D. simulans* can be explained by the genomic distribution of recombination-rate associated sequence motifs. We constructed the first high resolution recombination map for *D. simulans*, and searched for motifs linked with high recombination in both sister species. We identified five consensus motifs, present in either species. While the association between motif density and recombination is strong and positive in *D. melanogaster*, the results are equivocal in *D. simulans*. Despite the strong association in *D. melanogaster*, we do not find a decreasing density of these repeat motifs towards centromeres and telomeres. We conclude that the density of recombination-associated repeat motifs cannot explain the large-scale recombination landscape in *D. melanogaster*, nor the differences to *D. simulans*. The strong association seen for the sequence motifs in *D. melanogaster* likely reflects their impact influencing local differences in recombination rates along the genome.

## INTRODUCTION

Meiotic recombination rate variation impacts on multiple important biological processes in sexual eukaryotes. It is crucial for chromosomal segregation (John 2005; Roeder 1997), but is also itself a powerful factor influencing genome organisation and sequence variability (Aquadro, et al. 1994; True, et al. 1996). Meiotic recombination arises when a double-stranded break leads to crossing over between homologous chromatids (Bergerat, et al. 1997; Hughes, et al. 2018; Keeney, et al. 1997; Schwacha and Kleckner 1995; Szostak, et al. 1983). Higher rates of recombination break up genetic linkage and can increase the efficacy of natural selection (Charlesworth and Charlesworth 2010; Haddrill, et al. 2007) and so affect the evolution of numerous genomic features. The reduction of transposable element density (Charlesworth and Lapid 1992; Charlesworth, et al. 1994; Kofler, et al. 2012; Petrov, et al. 2011; Rizzon, et al. 2002) and the increased levels of DNA polymorphism (Aquadro, et al. 1994; Begun and Aquadro 1992; Begun, et al. 2007; Kulathinal, et al. 2008) in regions of high recombination are probably the clearest examples.

Yet while the eukaryotic meiotic machinery is generally highly conserved (Keeney 2001), rates of recombination have been observed to vary dramatically across species and populations, between individuals, and across sexes (Stapley, et al. 2017), apparently due to a combination of interacting environmental, epigenetic, and genetic factors (Detlefsen and Roberts 1921; Neel 1941; Parsons 1958; Stapley, et al. 2017; Stern 1926). Moreover, the distribution of meiotic recombination rates among and along chromosomes varies markedly across taxa (Choi and Henderson 2015; Hey 2004; Hunter, et al. 2016a; Lichten and Goldman 1995; Petes 2001; Stapley, et al. 2017). Large-scale recombination suppression is often observed towards centromeres, the so called “centromere effect” (Beadle 1932; Choulet, et al. 2014; Hughes, et al. 2018; Szauter 1984). Depending on the species, either suppression or enhancement of recombination has been observed towards the telomeres (Broman, et al. 1998; Chan, et al. 2012; Comeron, et al. 2012; Myers, et al. 2005). Heterochromatin, which is often associated with these regions, tends also to exhibit lower recombination rates than euchromatin (Baker 1958; Roberts 1965; Sturtevant and Beadle 1936; Szauter 1984; Termolino, et al. 2016). Yet, in addition to these large-scale features of recombination landscapes, fast-evolving (Jeffreys, et al. 2001) finer-scale variation can also be observed (Comeron, et al. 2012; Myers, et al. 2005).

It has been proposed that short sequence motifs are a key factor shaping the recombination landscape. For example, in humans a 13-mer, CCNCCNTNNCCNC motif is targeted by the PRDM9 protein (Billings, et al. 2013; Grey, et al. 2011; Myers, et al. 2010), via its zinc-finger array (Baudat, et al. 2010; Parvanov, et al. 2010), where it promotes histone methylation and meiotic crossover, reorganising the nucleosome around it and driving double stranded break formation (Baker, et al. 2014; Brick, et al. 2012; Mihola, et al. 2009; Pratto, et al. 2014). These highly localized recombination events in 500–2000bp sections of chromosome have been called recombination “hotspots” (Lam and Keeney 2014). They are observed in a multitude of species including yeast, mice, humans among many others (Lam and Keeney 2014).

Hotspots are, however, no universal feature of recombination landscapes, and are not observed in a range of species groups including *Caenorhabditis elegans* and *Drosophila* (Aquadro, et al. 2001; Chan, et al. 2012; Hey 2004; Manzano-Winkler, et al. 2013; Miller, et al. 2016; Nachman 2002; Smukowski Heil, et al. 2015). *Drosophila spp.*, exhibit a large heterogeneity in recombination across their chromosomes, as demonstrated in *D. persimilis* (Stevison and Noor 2010), *D. pseudoobscura* (Cirulli, et al. 2007; Kulathinal, et al. 2008), and *D. melanogaster* (Adrian, et al. 2016; Comeron, et al. 2012; Singh, et al. 2009). Still, *D. melanogaster* exhibits only a handful of mild “hotspots” relative to the ~30,000, often very strong hotspots observed in humans (International HapMap Consortium 2007). Instead the *D. melanogaster* recombination landscape is characterised by recombination “peaks” and “valleys” on a 5kb – 500kb scale (Adrian, et al. 2016; Chan, et al. 2012; Comeron, et al. 2012; Singh, et al. 2009) with which short “recombination motifs” are associated; as is also seen in *D. pseudoobscura, D. persimilis*, and other species (Adrian, et al. 2016; Chan, et al. 2012; Cirulli, et al. 2007; Comeron, et al. 2012; Heil and Noor 2012; Kulathinal, et al. 2008; Miller, et al. 2012; Singh, et al. 2009; Singh, et al. 2013; Stevison and Noor 2010). These motifs, which often reside in transcription-associated euchromatic regions (Comeron, et al. 2012; Petes 2001), are thought to increase the accessibility of DNA chromatin to double-stranded cleavage (Comeron, et al. 2012) and de-stabilize DNA sequences, potentially in a stress, environmental or epigenetically dependent manner (Hunter, et al. 2016b; Kohl and Singh 2018; Neel 1941; Petes 2001; Redfield 1966; Stern 1926).

*D. melanogaster, D. simulans*, and *D. mauritiana* are sister species which are ecologically and karyotypically similar (LEMEUNIER AND ASHBURNER 1976; TRUE *et al.* 1996), but differ dramatically in their recombination landscapes. While *D. melanogaster* exhibits a characteristic gradual decrease in recombination rate towards centromeres and to a lesser extent also telomeres, the recombination landscape in *D. simulans* and *D. mauritiana* is much flatter with a rather constant recombination rate almost to the end of the chromosome arm, where it drops very quickly (True, et al. 1996). Furthermore, these two species also have a higher recombination rate than *D. melanogaster* (True, et al. 1996), which has been attributed, in *D. mauritiana*, to the MEI-218 protein which has highly diverged between *D. melanogaster* and *D. mauritiana*, promoting recombination to a lesser extent in the former (Brand, et al. 2018).

Here, to test the hypothesis that differences in genome-wide motif distributions can explain the observed differences in recombination (Adrian, et al. 2016), we take a multi-step approach. First, we produce a high-resolution recombination map for *D. simulans*. Next, we run a motif discovery in each species and construct a consensus motif set. We confirm the clear differences in recombination landscapes between the two species, but find a similar set and distribution of recombination associated motifs in each. Our results suggest that recombination associated motifs cannot explain the large-scale differences in recombination landscapes between the two species but may have a significant impact on recombination on a local scale, in particular in *D. melanogaster*.

## MATERIAL AND METHODS

### Recombination Map

#### Recombination Map Production

A total of 202 isofemale lines were established from a natural *D. simulans* population in Tallahassee, Florida, USA in 2010 (Barghi, et al. 2017). From each of the 189 lines that were still alive in 2016, an individual male was selected and crossed with a virgin “reference” female from the M252 strain that was used to produce the *D. simulans* reference genome (Palmieri, et al. 2015). Paired-end libraries were generated for a single F1 female as described in Barghi, et al. (2017) and sequenced on an Illumina HiSeq XTEN to obtain an average sequence coverage of 30x. Single-nucleotide polymorphisms (SNPs) were called with FreeBayes (v1.1.0-46-g8d2b3a0, Garrison and Marth 2012), requiring a minimum sequencing coverage of 10x and a variant quality of at least 50. All SNPs that were polymorphic in the M252 reference strain were masked. Based on line-specific haplotype information, the genome-wide recombination map was estimated with LDJump (v0.1.4, Hermann, et al. 2018), specifying a segment size of 1kb, with an a = 0.05 and an Q = 0.04. We disabled LDJump’s segmentation analysis and worked with raw recombination rate estimates. Recombination rates were converted from r to units of cM/Mb by normalising them so as to have a genetic map length between a set of marker genes equivalent to that which has been previously reported (True, et al. 1996).

The resultant *D. simulans* recombination map was used in parallel with the *D. melanogaster* recombination map produced by Comeron, et al. (2012), downloaded from the *Drosophila melanogaster* Recombination Rate Calculator (Fiston-Lavier, et al. 2010).

#### Recombination Map Scaling

As the raw recombination map output by LDJump is noisy, we smoothed each recombination map at several scales. In *D. melanogaster*, the raw map (Comeron, et al. 2012) contained information on recombination rate at a 100kb resolution, in *D. simulans* raw information was generated at a 1kb scale. For smoothing, we used a moving median approach, using window sizes of 5, 25, 101, 501 and 2501 kb for *D. simulans*, and a 101, 501, 2501 kb for *D. melanogaster*, respectively. Advantages of the moving median as a smoothing method include low sensitivity to outliers, and a direct relationship to underling data, in the sense that only values present in the raw data set can be present in the smoothed set if the median is taken based on an odd number of input values, which in our case it always was. Because this approach is also computationally expensive, and prone to deleting map features when there are long runs of identical values, we investigated as an alternative approach, smoothing via LOESS local regression (Cleveland, et al. 1992), which produces qualitatively equivalent results (Figure S2). The smoothing scales chosen reflect those in Adrian, et al. (2016), relevant to potential motif explanatory power. The “correct” scale on which motifs may function is *a priori* unclear.

### DNA Motif Identification

#### Motif Discovery

For each species, we ran a genome-wide motif discovery using MEME (Bailey and Elkan 1994), from the MEME suite of motif-based sequence analysis tools (Bailey, et al. 2009, version 5.0.1pl, accessible at http://meme-suite.org; Bailey, et al. 2015), a software designed to detect DNA sequence motifs in genetic data. After dividing each of the five large chromosomes (X, 2L, 2R, 3L, 3R) into high- and low-recombining regions based on the chromosome median recombination rate, we used this software in the “differential enrichment” mode to detect motifs enriched in high-recombining areas of the genome. For *D. melanogaster*, we ran MEME on the release 5 reference genome (v. 5.36), for concordance with our recombination information from Comeron et al. (2012). For *D. simulans*, we used the M252 Madagascar reference genome (Palmieri, et al. 2015), to align with our recombination map. Motif discovery searches were run with species specific Markov Background Models, simple matrices of background base frequencies obtained using the MEME *fasta-get-model* command, for each reference genome in turn. The full procedure was repeated with all smoothed maps (Methods: *Recombination Map Production*). For completeness, a raw 1 kb window motif discovery run was also conducted for *D. simulans*. A similar search for motifs associated with lower recombination areas returned no results.

#### Motif Consensus Set

MEME motif discovery runs returned a set of 5, 4 and 3 motifs in *D. melanogaster* and 1, 2, 4, 1, 1 and 1 significant motifs in *D. simulans*, at the 101, 501, and 2501, and 1, 5, 25, 101, 501, and 2501 kb scales, respectively (SI.3, *E* ≤ 0.01). It was noticed that, while individually distinct, numerous motifs contained similar core patterns whilst varying, for example, only in repeat number. As such, we constructed a set of 5 consensus motifs that captured the core variation in all motifs significantly associated with increased recombination, across both species, and over all scales. This core set of motifs C1–5, was determined via a two-step method. First, we contrasted the motifs across each of our recombination map smoothing scales in both species, retaining only motifs that occurred in at least one scale with a minimum significance of *E* ≤ 0.01 in at least one species. Motifs were then simplified by allowing only the most likely base at each position, and motif lengths were fixed as the longest sequence length that could be represented in both species (as lengths were by tendency longer in *D. melanogaster*). This resulted in the following set of consensus motifs: C1=[A]_11_; C2=[GCA]_4_; C3=[CA]_6_; C4=[TA]_5_; C5=[G]_8_. We note that *D. melanogaster* made the dominant contribution to the consensus motifs, as the motifs in *D. simulans* were less significant than those observed in *D. melanogaster* (SI.3), and that the number of consensus motifs was informed by the data, and not decided *a priori*. As our consensus motifs turned out to be simplified versions of the most predictive motifs that were identified by Adrian et al. (2016), we quantitatively confirmed this similarity using the MEME Suite tool TomTom (Gupta, et al. 2007), under default parameters (see SI.4).

### Genome-Wide Motif Densities

#### Motif Locations

We converted the 5 consensus motifs into letter-probability matrices, to be used as input to FIMO, a MEME Suite tool designed to find genome-wide motif occurrences (Grant, et al. 2011). Matrices were compiled in a hard, and a softer, version; with the expected base given a probability of 1 and unexpected bases probabilities of 0, or the expected base a probability of 0.97, and unexpected bases a probability of 0.01. FIMO was then run for each species, taking the reference sequences and Markov Background Models as noted in Methods: *Motif Discovery*, and using parameter *max-stored-scores* = 50000000, and all others at default. Results of the hard and soft motif probability runs were qualitatively identical, so hard coded motif probabilities were used for follow-up analysis (soft runs not reported).

#### Motif Densities

FIMO output provides, per motif, the genomic locations (chromosome, start and stop position) at which a motif was found, as well as a *p*-value and a *q*-score (Benjamini and Hochberg 1995) per record, which show how well the motif was matched to the underlying reference sequence, both before and after correction for multiple testing (Benjamini and Hochberg 1995). To obtain genome-wide motif densities in each species, we calculated for each motif the sum of 1 – *q*, across a sliding window of 1 kb, where *q* refers to the per record *q*-score, such that per window motif densities are discounted in relation to the quality of the motif match, with higher quality matches counting more. A total, genome-wide count (of 1 – *q*) of each motif was also obtained from the raw FIMO output.

### Motif - Recombination Correlations and Models

#### Motif Density – Recombination Rate Correlations

To investigate the relationship between recombination rates and genome-wide abundances of individual motifs, we calculated the correlations between motif densities, binned at 1 kb, and corresponding recombination rates (cM/Mb), per motif, for *D. simulans* and *D. melanogaster*, respectively. As there was no clear *a priori* expectation for the genomic scale at which motifs would have most impact on recombination, the analysis was repeated for all smoothing scales noted in Methods: *Recombination Map Scaling* for *D. melanogaster* and *D. simulans* (and was repeated on the raw 1 kb scale in for *D. simulans*, not shown). Spearman’s roe, r, was used as a non-parametric estimator of the correlation between the test variables, and both the direction and significance of all correlations were extracted. To investigate the overall predictive power of motif densities, irrespective of chromosomal background, the analysis was repeated on the total genomic data, pooling across all of the 5 major chromosomes, with the analysis repeated per motif and species.

Finally, to test for explicit directional effects of each consensus motif on recombination, a linear regression model was fitted, per motif, species, scale, and chromosome, for the effect of motif density on local recombination rate, and repeated for the genome average.

A schematic representation of this analytic pipeline is presented in Figure 1. All statistical analyses were run in R, version 1.1383, using in house scripts (see SI.5).

**Fig. 1.**
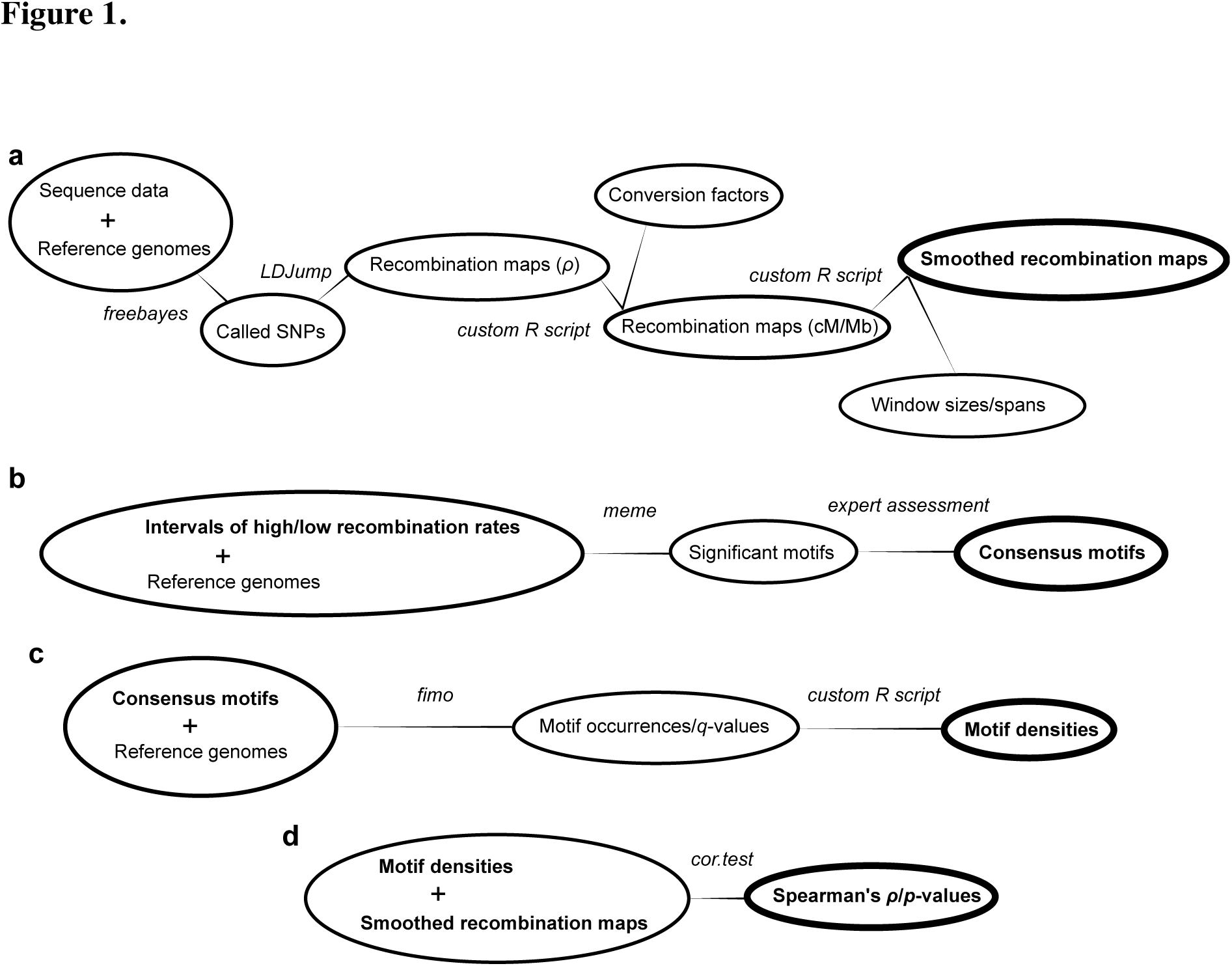
A schematic representation of the bioinformatic pipeline used. Ovals represent physical data sets, lines represent tools used to derive them; see the Methods for details.

## RESULTS

### Recombination rates in *D. simulans* are more uniform across chromosomes, than in *D. melanogaster*

We present the first high-resolution recombination map for *Drosophila simulans*, and contrast it to that of *D. melanogaster* (Comeron, et al. 2012). Across a range of smoothing parameters, the *D*. *simulans* recombination map is more uniform than that of *D*. *melanogaster* (Figures 2, 3). The level of recombination suppression is lower towards the centromere in *D. simulans*. As in *D. melanogaster*, the main broad-scale features of the *D. simulans* map hold across the full range of genomic scales, while finer resolution peaks and troughs become visible only at higher resolutions, at the 5 – 501 kb scale (Figure 3). The finer scale peaks (on a kb scale), as with the broader features (on a Mb scale), differ between these two sister *s*pecies, and persist across smoothing scales (Figures 2, 3).

**Fig. 2.**
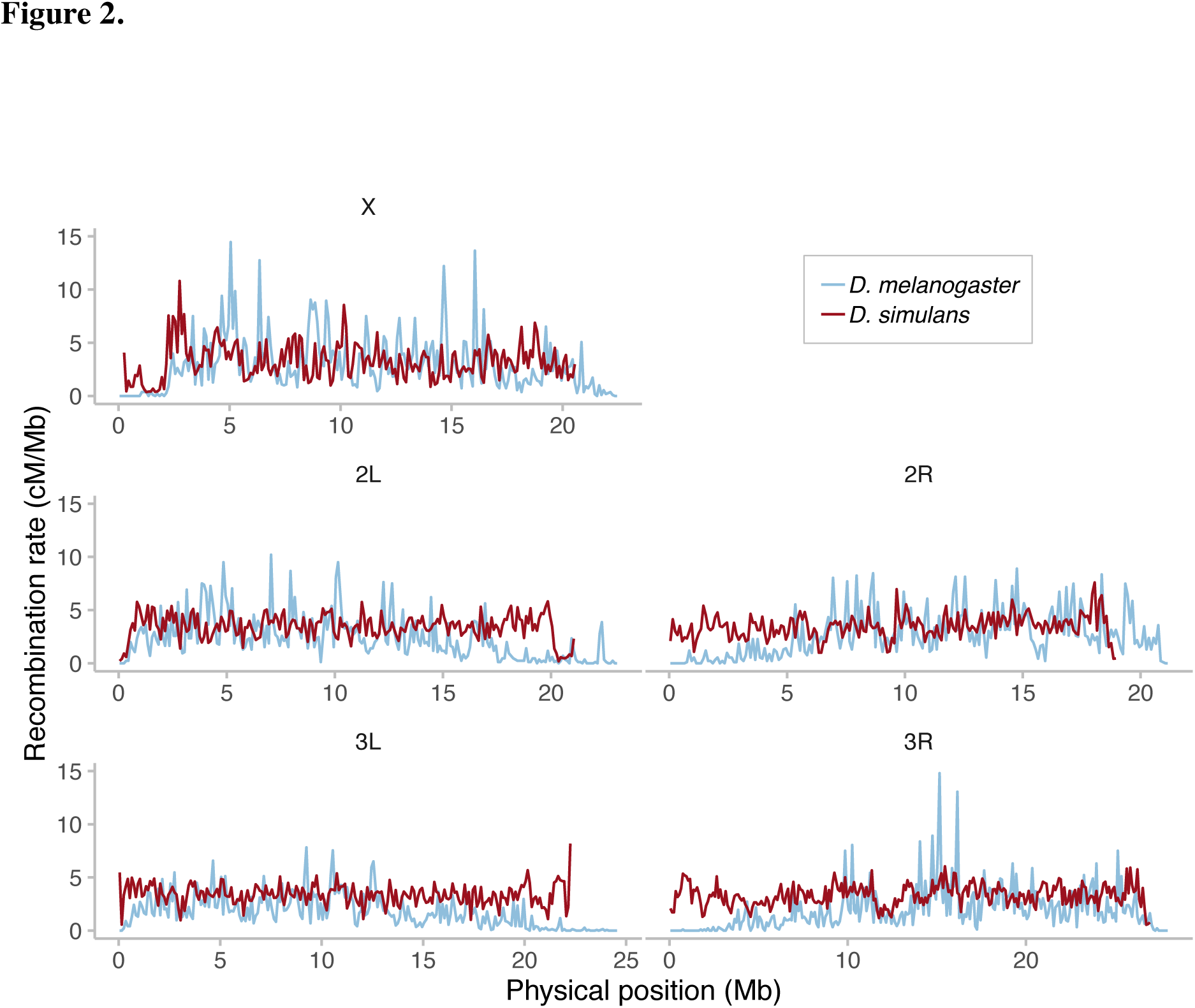
Recombination rates in *D. simulans* are more uniform across chromosomes than in *D. melanogaster*. Red lines show the recombination rate in *D. simulans* for each of the major chromosomes (name labels in top margin), smoothed at a 101 kb window size with a moving median. For comparison, blue lines show the recombination rate in *D. melanogaster* (with data taken from Comeron *et al*. 2012); Figure 3 for other resolutions.

**Fig. 3.**
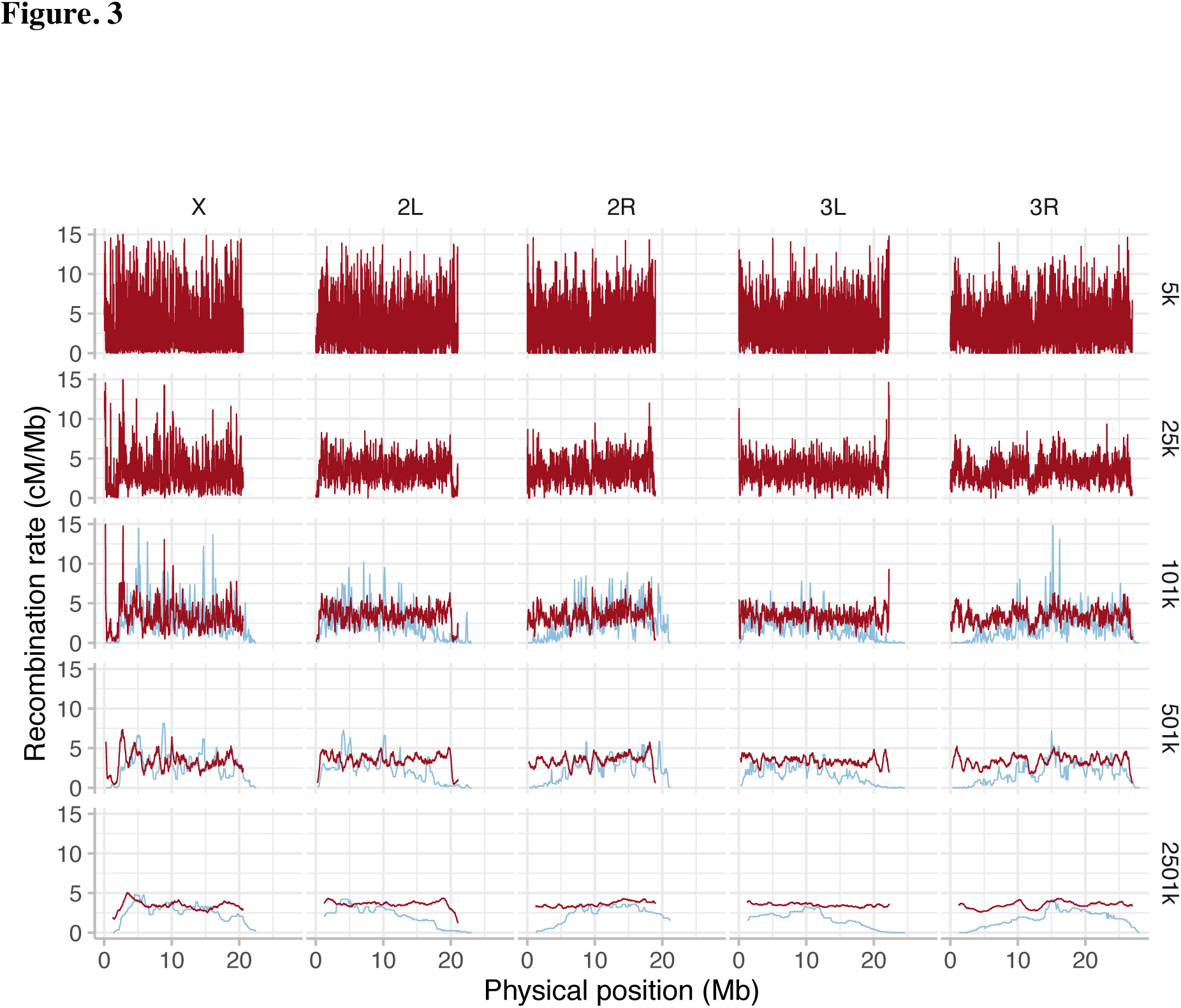
Recombination rates in *D. simulans* are more uniform across chromosomes than in *D. melanogaster*, at all smoothing scales. Red lines show the recombination rate in *D. simulans* for each of the major chromosomes (names in top margin), smoothed at 5 window sizes (right margin, in bp) with a moving median. For comparison, blue lines show the recombination rate in *D. melanogaster* (data taken from Comeron *et al*. 2012 at 101k; and smoothed at 501k and 2501k; with data not available at smaller resolutions).

### Motif density landscapes are similar in *D. simulans* and *D. melanogaster*

We identify 5 consensus motifs based on motifs recovered in each of the two species (Methods: *Motif Consensus Set*) and obtain their genome-wide densities. The consensus motifs were: C1=[A]_11_; C2=[GCA]_4_; C3=[CA]_6_; C4=[TA]_5_; C5=[G]_8_. Across all chromosomes and consensus motifs, motif density landscapes were similar in *D. melanogaster* and *D. simulans* (Figure 4). This was especially true for intermediate size landscape features, such as humps and wider valleys (e.g. motif C2 on X, 7.5 Mb position, or 2L at the 8 and 12 Mb positions, Figure 4). Therefore, motif density cannot explain the differences in the broad recombination landscape between both species.

**Fig. 4.**
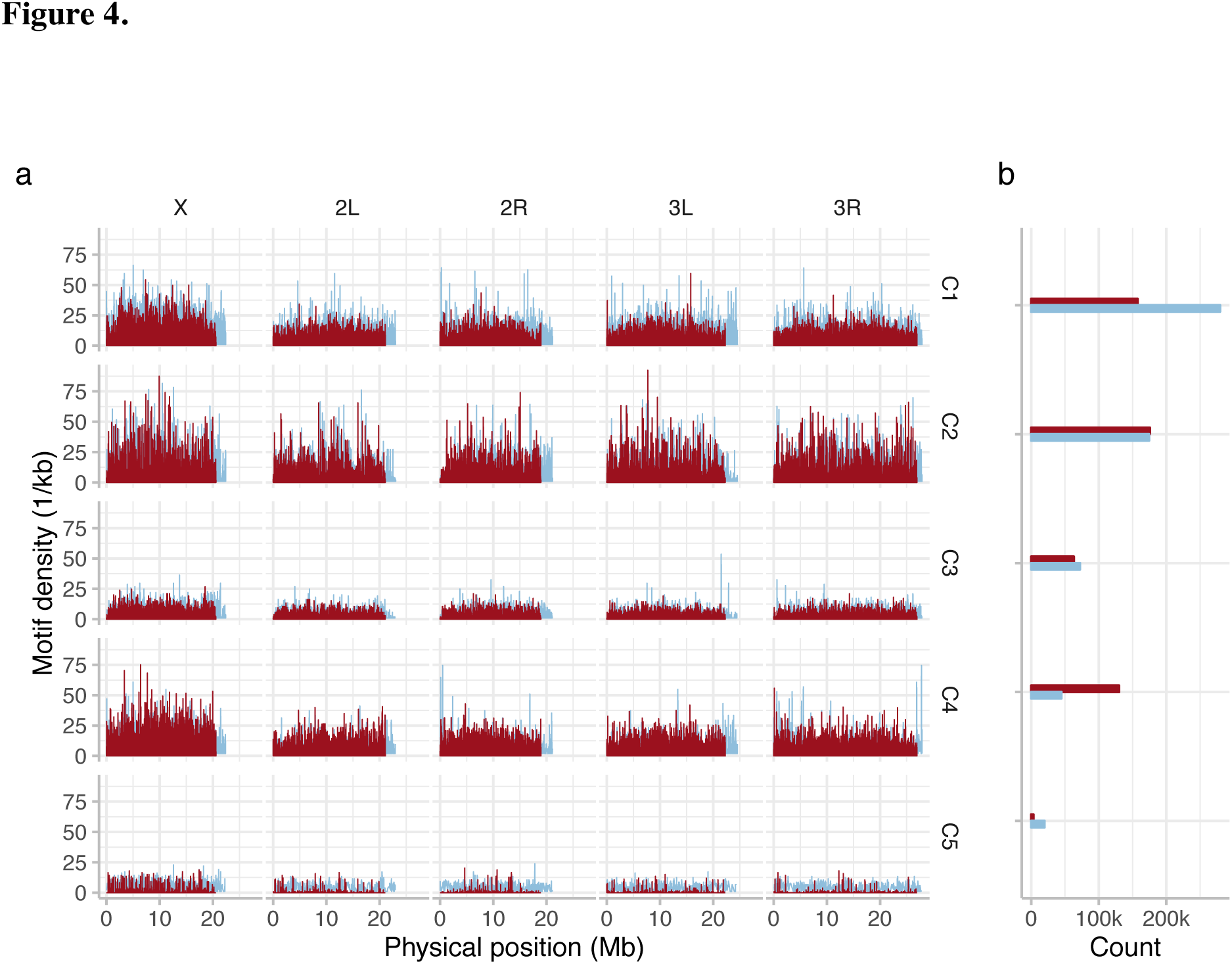
Motif densities are similar in *D. simulans* and *D. melanogaster*. (a) Red (*D. simulans*) and blue (*D. melanogaster*) lines show motif densities across major chromosomes (top margin) as reported by FIMO, with motif occurrences discounted by 1 – *q* (see main text) and binned into 1kb windows for each consensus motif (C1-5). (b) Total motif counts across all five large chromosomes. FIMO threshold: *p*-value of 1e-4 (default setting).

Finer resolution peaks and troughs varied more between species (e.g. motif C4 on X, 5–15 Mb position, Figure 4). Further, although the different motifs, C1–5, displayed similar broad patterns in each species – per chromosome and genome-wide – some species-specific patterns were seen. Motifs C1, [A]_11_ and C5, [G]_8_ were far less common in *D. simulans*, which had a lower total motif count, while the opposite was true for motif C4, [TA]_5_. Nonetheless, genome-wide motif distributions were similar in each species.

### Associations between motif densities and recombination rates are generally weaker and less significant in *D. simulans* than in *D. melanogaster*

We examined correlations between motif densities and recombination rates in each species, both per chromosome, and genome-wide, and at a range of genomic scales. A clear difference was observed between the species. In *D. melanogaster*, all but one correlation was positive, most were highly significant both genome-wide and per chromosome, and the correlation coefficients (Spearman’s r) were generally large; with a range of ~ 0.4 – 0.6 for the most associated motifs per chromosome (and genome-wide, Figure 5a). In contrast, the associations observed in *D. simulans* were heterogeneously positive or negative, had lower significances than those observed in *D. melanogaster*, and were in all cases weak; with a range of ~ 0.01 – 0.04 for the most associated motifs per chromosome (and genome-wide, Figure 5b). In both species, there was also variation in the importance of different motifs on different chromosomes (below). However, while in *D. melanogaster* the patterns of motif association held across all scales for each chromosome and genome-wide, in *D. simulans* there were occasional exceptions to this rule. For instance, on 2L, 2R, 3L, and genome-wide, the positive correlations for C1 and C4 switched direction at scales larger than 25 – 101 kb. Given that these correlations were very weak with low significance, we attribute these discrepancies stochastic noise, rather than biological signals. We finally note that motifs C1, C2, and C3 were the most associated with recombination across most major chromosomes in both species (though to a far lesser extent in *D. simulans*), but that an exception is observed for the X chromosome. Here, motif C2 had a very weak association with recombination rate in both species, and motif C4 instead had a high association, relative to its weak association on most autosomes in both species. Very similar observations were seen for the linear regressions (Figure S1), with more models being significant and positive for *D. melanogaster*.

**Fig. 5.**
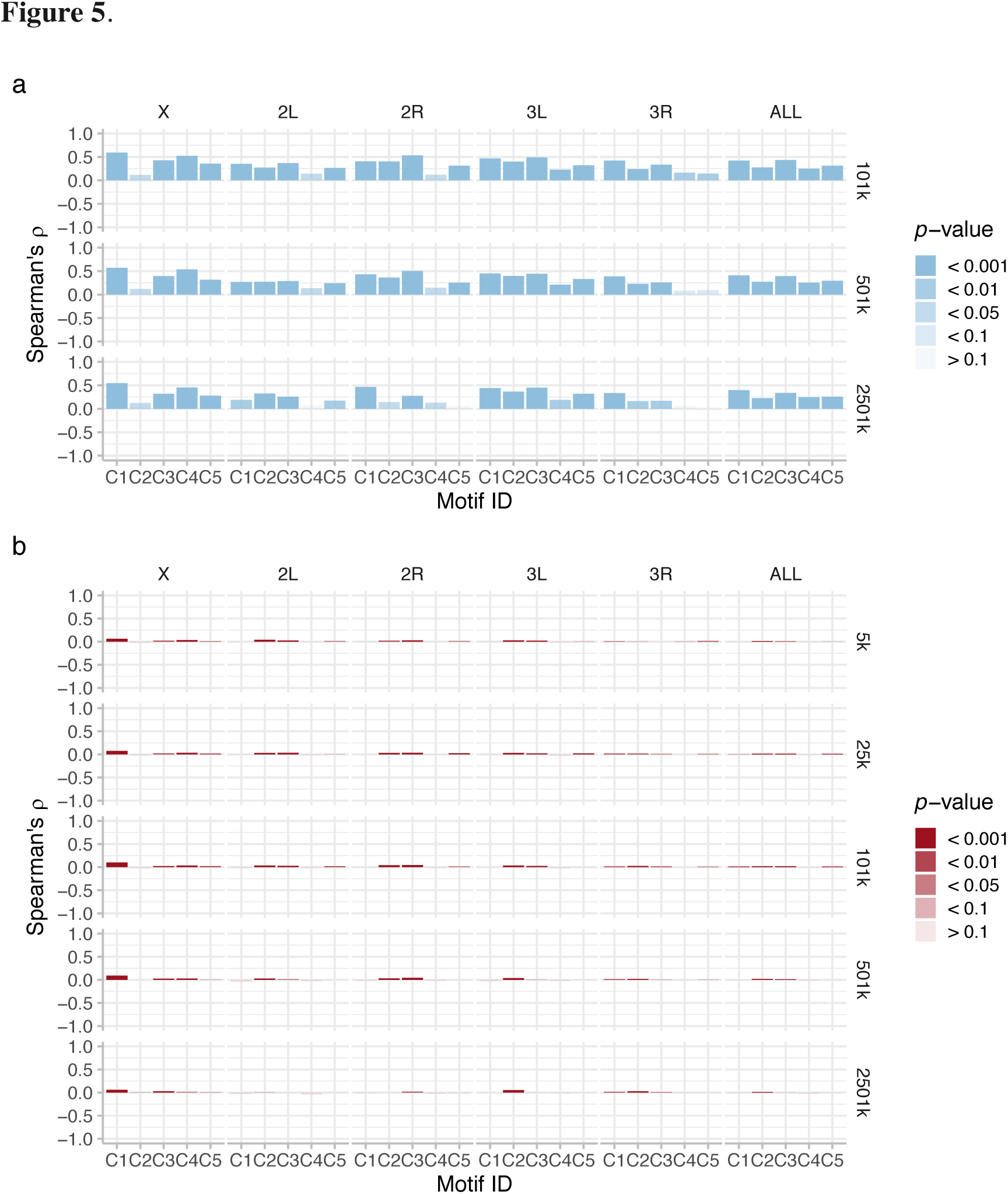
Associations between motif densities and recombination rates are generally weaker and less significant in *D. simulans* than in *D. melanogaster*. For (a) *D. melanogaster*, and (b) *D. simulans*, bars indicate Spearman's rho, r, (height) and the corresponding *p*-value (transparency), from tests of the correlation between motif densities (as shown in Figure. 4 for *D. simulans*, but re-binned for *D. melanogaster* to account for the resolution of the available data) and recombination rates, across individual chromosomes and for the five large chromosomes together (top margin), at all smoothing levels (see right margin).

## DISCUSSION

We present the first high resolution recombination map for *Drosophila simulans*, and a comparative analysis of recombination motifs and their association with recombination in two sister species, *D. melanogaster* and *D. simulans*. We tested the hypothesis that such motifs predict recombination rates within the *D. melanogaster* species subgroup.

Our *D. simulans* recombination map confirms the results of previous, lower resolution work in this species (Ohnishi and Voelker 1981, 1979; Stuktevanat 1929; True, et al. 1996). We find that the *D. simulans* recombination landscape is far flatter than in *D. melanogaster* (Figures 2, 3). While centromeric recombination suppression on the X, and to some extent on 2L and 3R, is observed in *D. simulans*, it is restricted to a small genomic region, whereas in *D. melanogaster* the recombination rate decreases only gradually over a much larger region in proximity to the centromeres (Comeron, et al. 2012). In *D. simulans*, a similarly sharp teleomeric suppression is also observed on 2L (and to some extent on X, 2R, 3L and 3R) at most smoothing scales (this pattern is less clear at 2501 kb). Unlike in *D. melanogaster*, overall recombination rates in *D. simulans* appear similar between X and the autosomes (Figures 2, 3) (Comeron, et al. 2012). We caution however, that recombination rate estimates from population polymorphism data are sensitive to demographic events and particular the ratio of X-chromosomal and autosomal variation differs widely between populations (Kauer, et al. 2002; Schöfl and Schlötterer 2004). In *D. simulans* and *D. melanogaster*, mid-to-large scale recombination features clearly persist over the 101, 501 and 2501 kb smoothed maps. In *D. simulans*, our high-resolution map shows that such features also persist down to the 25 and 5 kb scale (e.g. the dip on 3R at 12.5 Mb position, Figures 3). As with the centromeric differences however, mid-scale and narrower landscape features differ between the species, especially at the 101 and 501 kb resolutions. In short, at all genomic scales tested the two species differ dramatically in recombination rates, over broad- and finer-scale recombination features.

Direct implications from these differences in genetic maps are that linkage-disequilibrium should be both lower and less variable across the *D. simulans* chromosomes relative to those of *D. melanogaster*. It is important to keep in mind that the *D. simulans* reference genome includes less repetitive DNA at the centromeric and telomeric ends of the chromosomes, so a comparison of recombination rates in not possible at the extremes of these regions. Nonetheless, our results bolster the current understanding of *D. simulans* recombination as less heterogeneous than that of *D. melanogaster* (Comeron, et al. 2012; True, et al. 1996), and indicate that selection will be generally more efficient in *D. simulans*, as genes that are uncoupled by recombination selection may result in more distinct signals, in particular in Evolve and Resequence experiments (Barghi, et al. 2017; Kofler and Schlötterer 2014; Tobler, et al. 2014). Hence, adaptive evolutionary changes may occur more rapidly in *D. simulans*, all else being equal, because Hill-Robertson effects are reduced by the higher recombination (Hill and Robertson 1966).

Turning to the causes of this recombination variation, we ran a MEME motif search to identify short DNA sequence motifs associated with regions of higher than average recombination, repeating this search in both *D. melanogaster*, and *D. simulans*. The first point of note was that a larger number of motifs were returned in *D. melanogaster*, and that those in *D. simulans* were by tendency both shorter and showed a less significant association with recombination rate, with lower quality matches. Nonetheless, a generally similar set of motifs was recovered in each species, and across each recombination map smoothing scale investigated. In short, we obtained a subset of the *D. melanogaster* motifs in *D. simulans*; motifs C1, C5 and by trend, motifs C2 and C4, providing some confidence in the impact of these motifs on the recombination rate. The motif sharing between the two *Drosophila* species provides some evidence that recombination motifs may to some degree be universal across *Drosophila* species. This idea builds upon prior work, which has shown that there is some overlap in motifs between more distant *Drosophila* species, such as *D. pseudoobscura*, which exhibits CACAC (Cirulli, et al. 2007), CCCCACCCC and CCTCCCT motifs (Kulathinal, et al. 2008), and *D. persimilis*, which exhibits a CCNCCNTNNCCNC motif (Stevison and Noor 2010). This led Comeron, et al. (2012) to speculate that *Drosophila* has a stable set of recombination motifs of universal function, which they confirmed in part by showing that *D. melanogaster* also exhibit the CACAC and CCTCCCT motifs, though not the CCCCACCCC motif. Our study builds on this result, showing that a larger degree of motif overlap can be seen both when contrasting consensus motifs and when comparing between more closely related species, and that the [CA]_n_ motif is universal to all *Drosophila* species studied. However, it is immediately notable that no complex, multi-part motifs were recovered in our study.

The genome-wide distribution of motifs (Figure 4) revealed, somewhat surprisingly, that there are also clear parallels between the two species motif landscapes. Not only do motifs with higher density in *D. melanogaster* generally have a higher density in *D. simulans*, but the patterns of motif distribution genome-wide are also remarkably similar. For instance, a similar “hump” and “peak” can be observed at the 8 and 9 Mb positions of chromosomes X and 2L respectively, for motif C2, in both species, while a density “trough” can be seen at 15 Mb on chromosome 2L for this motif (Figure 4). Motifs C1, C3 and C4 likewise exhibit very limited differences between species, on all chromosomes (Figure 4), despite clear differences in recombination rates (Figure 3). A few differences do exist. Motif C1 is more common in *D. melanogaster*, even if the “landscape” is similar to *D. simulans*; Motif C5 is less common in *D. simulans*, and exhibits a distinct landscape on all autosomes; and, any narrow-scale features rarely overlap between species, mirroring patterns of distinct recombination peaks and similar landscapes seen in *D. melanogaster* populations (Chan, et al. 2012; Smukowski Heil, et al. 2015). Consequently, while it might be tempting to speculate that subtle differences in motif densities can explain the flatter recombination landscape of *D. simulans* and its unique recombination peak set, it is difficult to reconcile the distinctive patterns of recombination rate variation in the two species with their exceptionally similar motif density landscapes, that are almost identical between species, especially when focusing on the large-scale differences in centromeric and telomeric regions.

The similar motif density patterns between the two species cast doubt on the hypothesis that differences in motif distribution can account for differences in recombination variation in these species. If divergent motif densities really account for the species differences in recombination rates, how can we explain the lack of concordance between reduced recombination towards the centromeres in *D. melanogaster*, the lack of this reduction in *D. simulans*, and the similar motif distributions over these regions in both species? To investigate this observation quantitatively, we calculated Spearman’s r as an estimator of the correlation between genome-wide motif density and recombination rate (cM/Mb), for each motif, in each species, across a range of smoothing scales. This revealed a striking difference between the two species. In *D. melanogaster*, all associations (aside one) were positive, for all motifs at all scales tested, with low *P*-values observed in most cases (Figure 5). These results accord well with those of Adrian, et al. (2016), who found positive associations between motif densities and recombination rate in *D. melanogaster*, using a similar set of motifs. In contrast, the associations observed in *D. simulans* were far smaller, and far more heterogeneous across chromosomes and motifs (Figure 5). This observation was confirmed by our linear regression models, fitted to explicitly test the predictive power of each motif to explain recombination rate variation, which showed an almost identical pattern (Figure S1). The correlations and model fits were similar within each species across all smoothing scales, in that the level of correlation did not increase with higher or lower resolution recombination maps. The clear implication is that motif densities do not universally predict recombination rates across the *Drosophila* clade, and are in particular not responsible for the large-scale differences observed between our two species. It is therefore pertinent to ask what alternative mechanisms could explain such differences.

A strong candidate is the dicistronic meiosis gene *mei-217*/*mei-218* and its protein product, MEI-218 (Brand, et al. 2018), which is involved in the resolution of crossing over into double stranded breaks and recombination (Brand, et al. 2018). Divergent forms have recently been identified in *D. mauritiana* and *D. melanogaster*, species that diverged 0.6 – 0.9 Ma. Like *D. simulans*, *D. mauritiana* exhibits a higher and flatter recombination rate landscape than *D. melanogaster* (True, et al. 1996), with the difference especially pronounced in the centromeric and telomeric regions (True, et al. 1996), and with this pattern expressed to an even larger extent than is seen in *D. simulans* (True, et al. 1996). Intriguingly then, Brand, et al. (2018) also found a high divergence in DNA and protein structure in the *mei-217*/*mei-218* gene and MEI-218 protein between *D. mauritiana* and *D. melanogaster*. The *D. mauritiana* form was far more effective in promoting recombination, increasing recombination assurance and reducing crossover interference (Brand, et al. 2018). It explained a large portion of the variance in crossover rates between *D. mauritiana* and *D. melanogaster*, especially that in the centromeric and telomeric regions (Brand, et al. 2018), and so could be a primary mechanistic variant explaining the differences in recombination between *D. simulans* and *D. melanogaster*. The clear parallel differences between the recombination maps of *D. melanogaster* versus *D. mauritiana* and *D. melanogaster* versus *D. simulans* imply that *mei-217*/*mei-218* may be responsible for the heterogeneity in recombination landscape that we have observed.

What then might explain the clear correlations between motif density and recombination seen in *D. melanogaster*, but not *D. simulans*? A simple explanation is that motifs are responsible for variation in recombination rate on a local scale. Hence, the lower density in *D. simulans*, results also in less micro-scale variation in recombination rate. Alternatively, this pattern could be explained if the recombination motifs are recognised directly by cleavage proteins, similar to PRDM9, that differ in function or effectiveness between *D. simulans* and *D. melanogaster*. Recent evidence shows that a zinc-finger gene and protein of this type exists in *D. melanogaster* (Hunter, et al. 2016a). Yet, such proteins tend to bind to complex, rather than short-repeat motifs, making this explanation unlikely. Another possibility relates to chromatin structure, because short-repeat DNA recombination motifs are thought to play roles in loosening chromatin structure, increasing access for double-stranded break (DSB) inducing proteins (Adrian, et al. 2016, and references therein; Comeron, et al. 2012). This could account for micro-variation in recombination rates genome-wide between species, for instance because the motifs were generally shorter and so presumably less effective at chromatin loosening in *D. simulans*, genome-wide. Circumstantial evidence in favour of this hypothesis includes that in both species motif correlation patterns varied cross chromosomes – for instance, C4 was a good predictor only on X – suggesting that motifs can operate in a context dependent manner. Likewise, the removal of subcentromeric and subtelomeric region recombination data has been found not to alter correlational patterns in *D. melanogaster* (Adrian, et al. 2016), suggesting that if motifs densities explain some recombination rate genome wide, they cannot explain centromeric and telomeric differences.

In short, we present the hypothesis that while short-repeat DNA motifs may affect recombination at a micro-scale, genome-wide, for instance in relation to euchromatic structure context, they cannot explain the large differences in recombination landscape differences between species, especially in the centromeric and telomeric regions. This variation seems far more likely to be explained by a mechanism such as *mei-217*/*mei-218*.

## SUPPLEMENTARY MATERIAL

Supplementary Figures 1-2, and supplements 3-5 are available at [insert location here]. All data generated and used in this study are available (see Data Accessibility below for details).

## DATA ACCESSIBILITY

Raw sequence reads for the 189 isofemale line haplotypes are available to download at the European Nucleotide Archive (ENA) under the accession numbers: [[added on acceptance]]. Phased haplotypes are available from Dryad via accession numbers: [[added on acceptance]]. Finally, CSV files for the *D. simulans* recombination map, at each resolution, are available for download from Dryad via accession numbers [[added on acceptance]].

## ACKNOWLEDGMENTS

The authors thank all members of the Institute of Population Genetics for discussion and support on this project. This work was supported by the European Research Council (ERC) grant “ArchAdapt” to CS, an Austrian Academy of Sciences DOC fellowship to TT, and the Austrian Science Fund (FWF, W1225).

## AUTHOR CONTRIBUTIONS

RM, JMH and CS conceived the study and interpreted the results. TT produced the recombination map. VN generated the NGS libraries. RM performed the analysis, with input from JMH. JMH and C.S. wrote the manuscript with input from all authors.

## SI. Legends

**Fig. S1.** Linear models predict some of the variance in recombination rate in *D. melanogaster*, but not in *D. simulans*. Scatterplot of recombination rate vs. motif density for (a) *D. melanogaster* and (b) *D. simulans* (species also indicated by blue and red colour, respectively). Gray lines represent single-motif linear model fits, inset numbers the corresponding *r*^2^ values, and appended asterisks indicate the *p*-values of the model fits at * < 0.05, ** < 0.01, *** < 0.001. For purposes of this comparison only, smoothed *D. simulans* data at 101k is shown here, with the same resolution of the *D. melanogaster* data. The recognisable correlation features in (b) are unaffected by this downsampling step (not shown).

**Fig. S2.** Loess-smoothed recombination maps. Red lines show the recombination rate in *D. simulans* for each of the major chromosomes (name labels in top margin), smoothed at 4 window sizes (see right margin, in bp) with the LOESS span parameter. LOESS span parameters correspond to 25, 101, 501, and 2501 kb, as span parameters equivalent to 5 kb can’t be implemented. For comparison, blue lines show the recombination rate in *D. melanogaster* (with data taken from Comeron *et al*. 2012 at 101k; and then smoothed at 501k and 2501k; with data not available at smaller resolutions).

**S3.** MEME motif discovery output for each *Drosophila* species at each genomic resolution.

**S4.** TomTom contrast of motifs from Adrian et al. (2016) to our set of 5 consensus motifs.

**S5.** R-Markdown document with script to reproduce our results [on acceptance of the article].

